# Guide-specific loss of efficiency and off-target reduction with Cas9 variants

**DOI:** 10.1101/2023.03.16.532856

**Authors:** Liang Zhang, Wei He, Rongjie Fu, Han Xu

## Abstract

High-fidelity Cas9 variants have been developed to reduce the off-target effects of CRISPR systems at a cost of efficiency loss. To systematically evaluate the efficiency and off-target tolerance of Cas9 variants in complex with different single guide RNAs (sgRNAs), we applied high-throughput viability screens and a synthetic paired sgRNA-target system to assess thousands of sgRNAs in combination with two high-fidelity Cas9 variants HiFi and LZ3. Comparing these variants against WT SpCas9, we found that ~20% of sgRNAs are associated with a significant loss of efficiency when complexed with either HiFi or LZ3. The loss of efficiency is dependent on the sequence context in the seed region of sgRNAs, as well as at positions 15-18 in the non-seed region that interacts with the REC3 domain of Cas9, suggesting that the variant-specific mutations in REC3 domain account for the loss of efficiency. We also observed various degrees of sequence-dependent off-target reduction when different sgRNAs are used in combination with the variants. Given these observations, we developed GuideVar, a transfer-learning-based computational framework for the prediction of on-target efficiency and off-target effect with high-fidelity variants. GuideVar facilitates the prioritization of sgRNAs in the applications with HiFi and LZ3, as demonstrated by the improvement of signal-to-noise ratios in high-throughput viability screens using these high-fidelity variants.

## Introduction

CRISPR-Cas9 technology is a widely used tool for genome editing and is currently being tested in clinical trials for therapeutic applications (1–5). Many applications of this technology utilize wild-type *Streptococcus pyogenes* Cas9 (WT SpCas9) owing to its high on-target activity. However, the off-target effects of the WT SpCas9 have raised critical concerns in scientific and clinical applications (6–9). To address this issue, high-fidelity Cas9 variants have been developed, including eSpCas9 (10), SpCas9-HF1 (11), HypaCas9 (12), evoCas9 (13), xCas9 (14), Sniper-Cas9 (15) and HiFi (16), etc. Despite the improved specificity of these engineered or evolved Cas9 variants, they often sacrifice the editing efficiency compared to WT SpCas9 (17). Recently, two promising Cas9 variants, HiFi and LZ3, have been reported to be associated with enhanced specificity while maintaining comparable on-target activity to WT SpCas9 (16,18). However, the cleavage activities of these high-fidelity variants were only tested in complex with a limited number of sgRNAs, and it is unclear if the improvements made by the variants are general for all sgRNAs or are specific to a subset of sgRNAs.

The efficiency and off-target tolerance of CRISPR-Cas9 system are not only affected by the Cas9 protein, but also the sequence context of the sgRNA (19–21). We and others have proposed sets of sequence rules for the prediction and optimization of sgRNAs with WT SpCas9 (7,19,22,23). Some of the rules, such as a guide-intrinsic mismatch tolerance (GMT) that can be explained by a kinetic model, are generally applicable to different Cas9 variants (24). However, structural analysis has shown that the mutations introduced to the Cas9 variants significantly alter the interacting patterns between the Cas9 protein and the RNA/DNA heteroduplex (12). Therefore, it is expected that the sequence determinants underlying the sensitivity and specificity of Cas9 variants could be divergent from the rules derived for WT SpCas9 (21).

To systematically explore the variant-specific sequence rules that determine the editing efficiency of Cas9 variants, we applied high-throughput viability screens to evaluate the efficiency of ~24,000 sgRNAs complexed with HiFi, LZ3, or WT SpCas9. Moreover, we measured the off-target tolerance of Cas9 variants in context of 328 sgRNAs and 1,753 target sequences using a synthetic paired sgRNA-target system. These data allowed a comprehensive modelling of efficiency and off-target effect of the variants, which further led to the development of a new machine-learning framework for sgRNA design in application with the high-fidelity Cas9 variants.

## Results

### Guide-specific loss of efficiency revealed by high-throughput knockout screens with Cas9 variants HiFi and LZ3

To systematically compare the editing efficiency between WT SpCas9 and Cas9 variants, we performed high-throughput viability screens in cell lines with stable expression of WT SpCas9, HiFi, or LZ3. A colorectal adenocarcinoma cell line DLD-1 and a small cell lung cancer cell line H2171 were individually engineered with WT SpCas9 or the two Cas9 variants by lentiviral transduction. Western blot confirmed equivalent expression of the three types of Cas9 and effective gene knockouts in all 6 engineered cell lines (Figure S1). The procedures of high-throughput viability screens are demonstrated in Figure 1A and detailed in Methods. Here, we used two independent lentiviral plasmid libraries, each containing ~12,000 sgRNAs (Figure S2). The first library, named EpiC, contains sgRNAs targeting ~1,000 epigenetic regulators and cancer-related genes, as well as control sgRNAs targeting core essential and nonessential genes (25). The second is a tiling-sgRNA library that contains all sgRNAs targeting the exons of 26 essential transcription factors required for the proliferation and survival of DLD-1 cells. Of the 6 engineered cell lines, the DLD-1 cells were transduced with either the EpiC or Tiling libraries, and the H2171 cells were transduced with the EpiC library. After ~20-day selection, the sgRNA sequences were amplified, sequenced, and counted for abundance relative to plasmid DNA.

**Figure 1.**
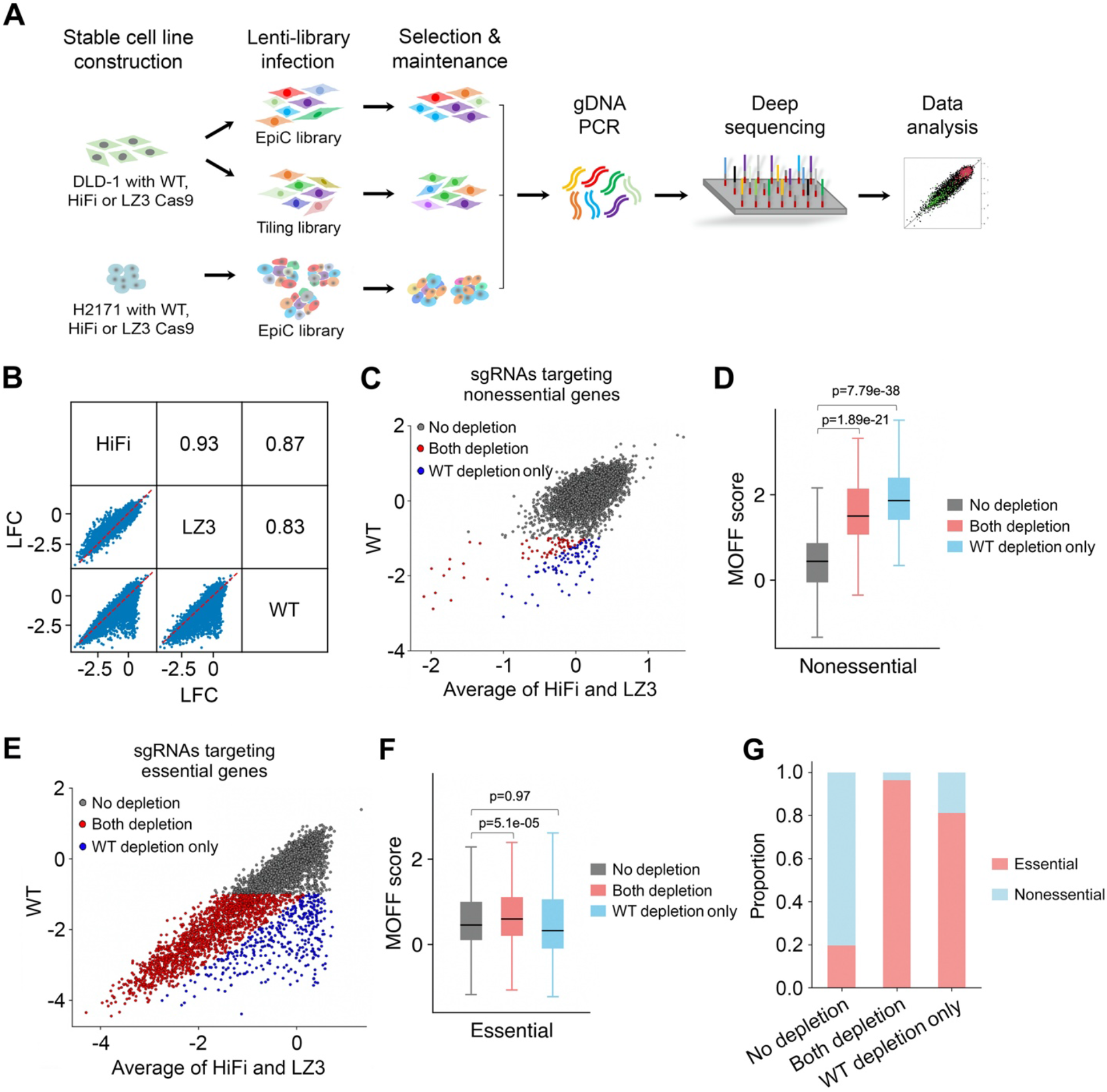
Performance of high-fidelity Cas9 variants in high-throughput CRISPR/Cas9 viability screens. **(A)** Experimental procedures of CRISPR screens using WT SpCas9, HiFi and LZ3 in combination with different sgRNA libraries in DLD-1 and H2171 cells. **(B)** Pairwise comparison of the Log Fold Changes (LFCs) of sgRNAs among the screens using WT SpCas9, HiFi or LZ3 in DLD-1 cells with EpiC library. The consistency of dropout effects was measured in Pearson correlation. **(C)** Scatter plot comparing the LFCs of sgRNAs targeting nonessential genes in WT SpCas9 and variant screens in DLD-1 cells with EpiC library. The sgRNAs were classified into three categories: “No depletion” (grey dots) and “Both depletion” (red dots) for both WT SpCas9 and the variants, and “WT depletion only” (blue dots). **(D)** Boxplot showing the predicted off-target effects (MOFF score) of the sgRNAs targeting nonessential genes in each category in (C). **(E)** Scatter plot comparing the LFCs of sgRNAs targeting essential genes in WT SpCas9 and variant screens in DLD-1 cells with EpiC library. **(F)** Boxplot showing the predicted off-target effects of the sgRNAs targeting essential genes in each category in (E). **(G)** The proportion of the sgRNAs targeting nonessential or essential genes, for each sgRNA category in (C) and (E).

We first tested the consistency of dropout effects among the screens with WT SpCas9, HiFi, and LZ3, based on pairwise Pearson correlations of the Log Fold Changes (LFCs) of sgRNA abundance (Figure 1B and Supplementary Table S1). In DLD-1 cells transduced with the EpiC library, HiFi and LZ3 showed a high correlation (r=0.93), whereas the correlations decreased when comparing HiFi and LZ3 against WT SpCas9 (r=0.87 and 0.83, respectively). Upon further examination, we found that a fraction of sgRNAs was associated with significantly less LFCs when HiFi and LZ3 were used, as compared to the screen with WT SpCas9 (Figure 1B). This observation was reproducible in DLD-1 cells transduced with the Tiling library and in H2171 cells transduced with the EpiC library (Figure S3A and B, Supplementary Table S2 and S3). Thus, the correlation analysis suggests a guide-specific variant-dependent cell dropout effect.

We reason that two factors may account for the decreased dropout effects with Cas9 variants. First, the variants reduce the off-target effect which could lead to a defect in cell viability. Second, some sgRNAs can be associated with loss of efficiency when targeting essential genes using the variants. To dissect these two factors, we examined the sgRNAs in the EpiC library that target nonessential and essential genes individually. Considering the high correlation between HiFi and LZ3, we averaged the LFCs of sgRNA abundance corresponding to the two variants in DLD-1 cells for further analysis. For the sgRNAs targeting nonessential genes, 151 (1.8%) out of 8,224 showed high dropout effect (LFC<-1) in the WT SpCas9 screen (Figure 1C). As expected, the dropped-out sgRNAs are associated with high off-targeting potential measured by the MOFF score (22), indicating that a small fraction of sgRNAs leads to viability defect of cells independent on the function of their target genes due to the off-target effects (Figure 1D). Among them, 88 (58.3%) showed decreased dropout effects (>2-fold difference in sgRNA abundance between the variants and WT SpCas9 screens) in HiFi and LZ3 screens, suggesting that the majority of the off-target effects were reduced by the variants. For the sgRNAs targeting essential genes, 2,021 (50.7%) of 3,990 showed dropout effects (LFC<-1) in the WT SpCas9 screen. Among them, 379 (18.8%) sgRNAs are associated with decreased dropout effects (>2-fold difference) in the variant screens compared to the WT SpCas9 screen (Figure 1E, blue dots). Interestingly, those sgRNAs depleted only in the WT SpCas9 screen are associated with similar MOFF scores to those without dropout effect, suggesting that the difference in dropout effects is mainly caused by the loss of efficiency of the sgRNAs in variant screens instead of the reduction of off-target effects (Figure 1F). Indeed, among all the sgRNAs in the EpiC library that were depleted only in the WT SpCas9 screen, nearly 80% target essential genes where on-target functional perturbations lead to phenotypic changes (Figure 1G and Supplementary Figure S3E). We observed similar results to the EpiC library in H2171 cells, and to the Tiling library in DLD-1 cells (Supplementary Figure S3C and D). Taken together, these lines of evidence indicate that sgRNA-specific loss of efficiency is the major causal factor that accounts for the difference in dropout effects between the screens with Cas9 variants and that WT SpCas9.

### Sequence features that contribute to the loss of efficiency with Cas9 variants

Next, we asked if the nucleotide sequence context of sgRNAs contributes to the loss of efficiency in Cas9 variant screens. To answer this question, we combined the sgRNAs targeting essential genes in the EpiC and Tiling libraries and categorized them into three groups: “Inefficient” (not efficient in either WT SpCas9 or Cas9 variant screens), “WT efficient only” (efficient only in WT SpCas9 screen) and “Both efficient” (efficient in both WT SpCas9 and Cas9 variant screens). We then computed the log odds ratios of spacer nucleotide frequency between the “Both efficient” and “WT efficient only” groups. We observed highly consistent sequence features between HiFi and LZ3 (Figure 2A and B, Supplementary Figure S4). We further extended the analysis to an orthogonal dataset generated through direct measurement of indel frequency induced by WT SpCas9 or HF1, another high-fidelity Cas9 variant (26). Despite different types of Cas9 variants and experimental settings, many sequence features are reproducible among HiFi, LZ3 and HF1 (Figure 2C and Supplementary Figure S4), suggesting a common mechanism underlying the sequence-dependent loss of efficiency among the three variants. To determine the importance of nucleotide positions, we calculated Kullback-Leibler (KL) divergence that measures the difference of nucleotide distribution at each position between the “Both efficient” and “WT efficient only” groups. We found that sequences in three regions significantly contribute to the loss of efficiency: i) positions 2-5 in the seed region proximal to the PAM, ii) positions 7-11, excluding position 9, at the junction of the seed and non-seed regions, and iii) positions 15-18 in the non-seed region (Figure 2D).

**Figure 2.**
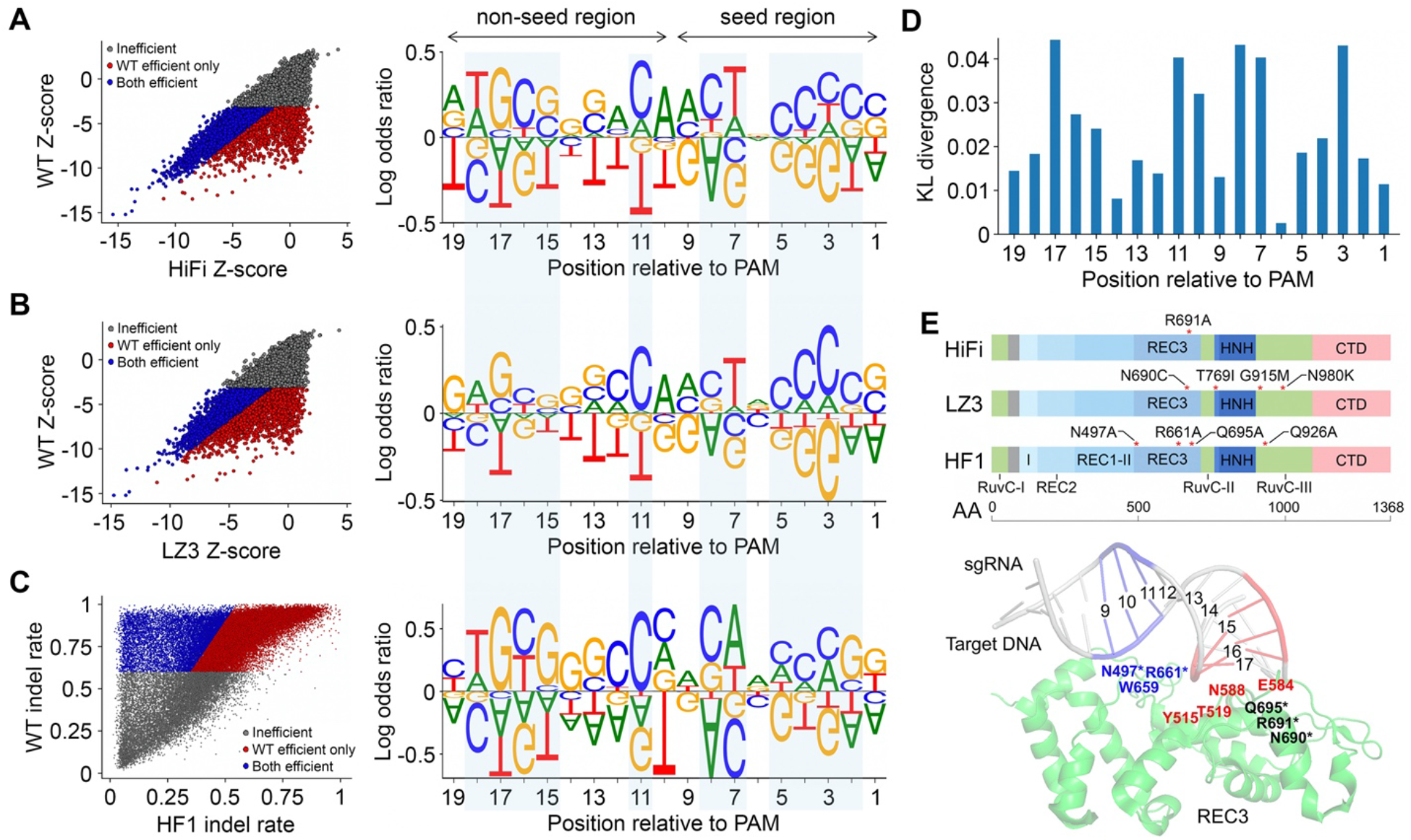
Sequence features associated with efficiency loss with Cas9 variants. **(A, B)** Left: scatter plot showing the categorization of sgRNAs into three groups: “Inefficient” (grey), “Both efficient” (blue), and “WT efficient only” (red), based on the screens with (A) HiFi and (B) LZ3. Right: the log odds ratios of nucleotide frequency between “Both efficient” and “WT efficient only” groups. The data were retrieved by combining all sgRNAs targeting essential genes in the EpiC and Tiling libraries. **(C)** Left: scatter plot showing the three sgRNA groups defined based on the indel rates in a published dataset using WT SpCas9 and the Cas9 variant HF1 (26). Right: the log odds ratios of nucleotide frequency between the groups of “Both efficient” and “WT efficient only”. **(D)** The KL divergence representing the significance of nucleotide difference between the “WT efficient only” and “Both efficient” groups at each position of the spacer. The KL divergence was averaged among the variants HiFi, LZ3, and HF1. **(E)** Upper: the point mutations (red asterisks) introduced to HiFi, LZ3 and HF1. Bottom: the structural representation of local interactions between the RNA/DNA heteroduplex and REC3 domain of WT SpCas9 (PDB: 7S4U). The positions 9-11 interacting with residues are marked in blue, the positions 15-17 interacting with residues are marked in red, and the residues mutated in any of the three Cas9 variants are marked with asterisks.

Previous reports indicated that sgRNA efficiency with WT SpCas9 is highly associated with the nucleotide sequence in the seed region (3,7,27). With regard to the variants, however, our results showed that the sequence context in the non-seed region also contributes to the variant-specific editing efficiency. We hypothesized that the observed sequence features in the non-seed region are associated with the structural alteration mediated by the mutations introduced to the variants. Comparing the mutations in HiFi, LZ3 and HF1, we found that all three variants harbor mutations in REC3, a non-catalytic domain that targets complementarity and governs the HNH nuclease to regulate catalytic competence (Figure 2E, upper) (11,16,18,28). Interestingly, the structural analysis revealed that positions 9-11 and 15-17, but not 12-14, interact with the REC3 domain of WT SpCas9 protein (Figure 2E, bottom) (29). Of note, it has been reported that mutations in REC3 can disrupt the interaction between REC3 and the RNA/DNA heteroduplex, while the interaction is required for the catalytic function (12,28,30,31). Therefore, our results support a model in which specific spacer sequences at positions 15-18 and/or 9-11 mediate the activity of Cas9 variants via conformational regulation of REC3-RNA/DNA interaction.

### Sequence-dependent off-target tolerance of Cas9 variants

Despite Cas9 variants being able to greatly reduce off-target effects compared to WT SpCas9, some sgRNAs targeting nonessential genes showed significant dropout effects in our viability screens when HiFi and LZ3 were used, corresponding to high off-target tolerance (Figure 1C and D). To systematically investigate the sequence-specific off-target tolerance of the Cas9 variants, we evaluated the off-target effects of 328 sgRNAs using a synthetic paired sgRNA-target system that allows high-throughput measurement of on-target and off-target indel rates in parallel (Figure 3A) (22). For each sgRNA, we designed 7 off-target sequences, including 3 targets with 1 mismatch (1-MM), 3 with 2 mismatches (2-MM), and 1 with 3 mismatches (3-MM). In consistency with the viability screens (Figure 1B, Supplementary Figure S3A and B), the on-target indel rates of HiFi and LZ3 showed a high correlation, whereas the correlations between the variants and WT SpCas9 were much lower (Supplementary Figure S5). In general, the off-target effects, as measured by off-on ratios, were significantly lower for the two Cas9 variants compared to WT SpCas9 (Figure 3B and Supplementary Table S4), confirming the overall reduced off-target tolerance of Cas9 variants. The off-target rates of HiFi and LZ3 were also highly correlated, in comparison to the reduced correlation between the variants and WT SpCas9 (Figure 3C).

**Figure 3.**
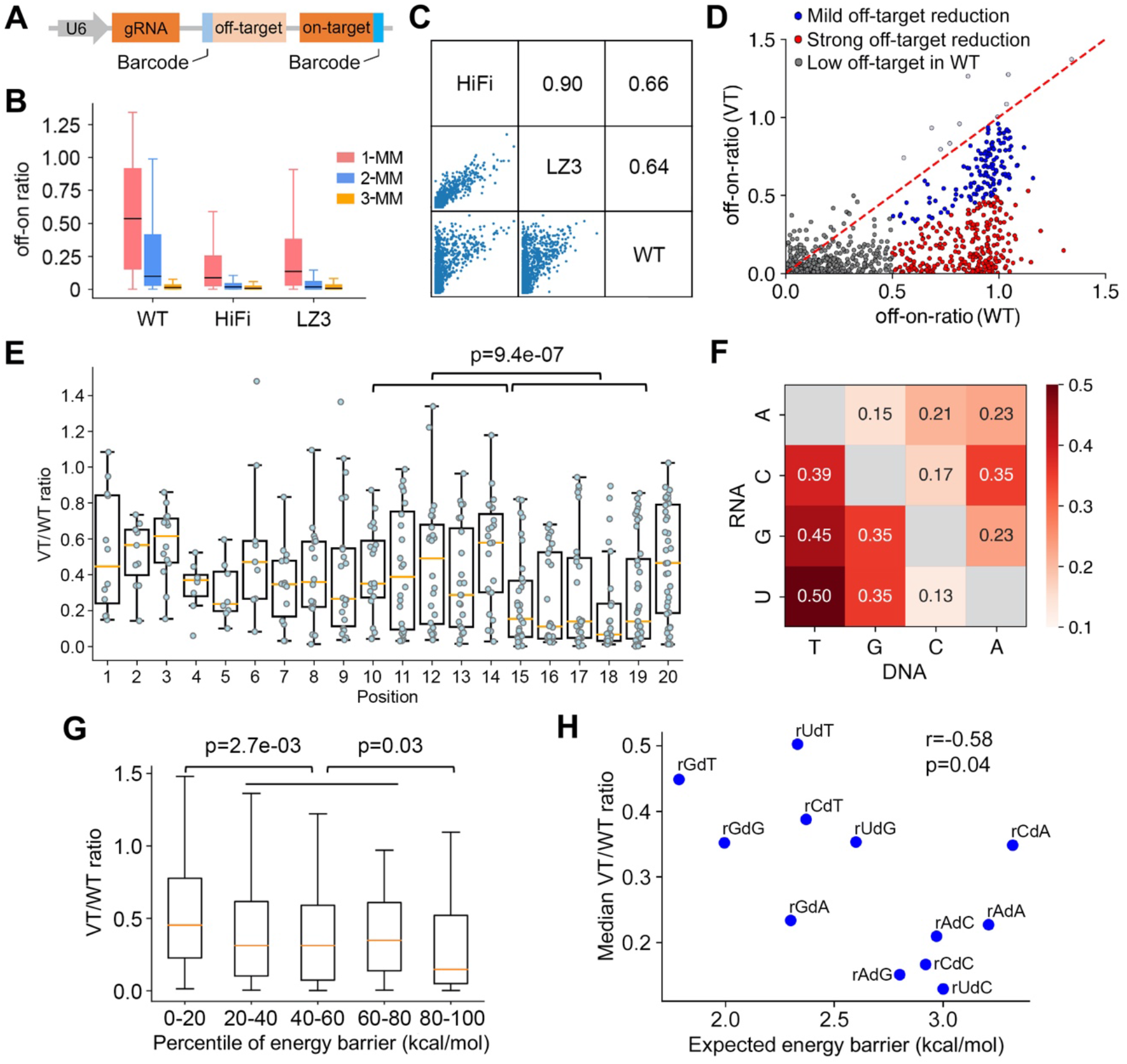
Sequence-specific off-target tolerance of Cas9 variants. **(A)** Schematic plot of the synthetic dual-target system for the evaluation of off-target effects for a specific sgRNA-target pair. **(B)** Boxplot comparing the off-on ratios of sgRNA-target pairs harboring 1-, 2-, or 3-mismatch (MM) among the screens with WT SpCas9, HiFi and LZ3. **(C)** Pairwise comparison of the off-target rates of sgRNA-target pairs among the screens with WT SpCas9, HiFi and LZ3. The consistency of off-target effects is measured in Pearson correlation. **(D)** Scatter plot comparing the off-on ratios of all the 1-MM sgRNA-target pairs between WT SpCas9 and the Cas9 variants (VT: average off-on ratio of HiFi and LZ3). The sgRNA-target pairs are assigned to three groups based on the off-target effects with WT SpCas9 and the degree of off-target reduction by the variants. **(E)** Boxplot showing the variant-mediated off-target reduction at each mismatch position measured as the ratio of off-target effects (off-on ratio) between the variants and WT SpCas9 (VT/WT ratio). The *p*-value is calculated using the two-tailed Manny-Whitney *U* test. **(F)** The median VT/WT ratios corresponding to different mismatch contexts between the sgRNA and the target DNA. **(G)** Comparison of VT/WT ratios at indicated percentile of expected energy barriers caused by the mismatches during R-loop formation. **(H)** Scatter plot showing the association between the energy barrier caused by a specific mismatch type and the median VT/WT ratio for that mismatch type. rXdY refers to mismatch type where the sgRNA sequence is X and the target DNA sequence is Y.

Upon further examination, we found that 317 sgRNA-target pairs were associated with different degrees of reduction of off-target effects when the variants were used (Figure 3D). To quantify the reduction of off-target effects by the variants, we computed the ratio of off-target effects between the variants and WT SpCas9 (VT/WT ratio) for 317 1-MM sgRNA-target pairs that are associated with strong off-target effects with WT SpCas9 (off-on ratio >0.5). We then asked if the position of mismatch accounts for the variation of VT/WT ratio. Interestingly, we observed significantly lower VT/WT ratios when a mismatch occurs at positions 15-18 of the RNA-DNA duplex that interact with the REC3 domain of Cas9 (Figure 3E). This observation suggests that the variant-specific mutations of REC3 and the mismatches at positions 15-18 can be synergistic in blocking Cas9-mediated DNA cleavage. In addition to the mismatch positions, we also found that the nucleotide contexts of RNA-DNA mismatches are associated with different degrees of VT/WT ratios (Figure 3F). In previous studies, we and others have reported that an RNA-DNA mismatch leads to an energy barrier during R-loop progression, where the quantity of energy barrier is predictable from the context of the mismatch and its adjacent nucleotides (22,32–34). Therefore, we computed the expected energy barrier based on the sequences of the 317 sgRNA-target pairs and found that those pairs with greater energy barriers are associated with stronger reduction of off-target effect by the variants (Figure 3G). Among the 12 single mismatch types, rUdC (i.e., uracil in RNA and cytosine in DNA), rCdC, rAdG, rAdC, and rAdA are associated with greater energy barriers and smaller VT/WT ratios (Figure 3H), suggesting that the variants are more effective in overcoming the off-target effects at the genomic off-target sites harboring these mismatches. Taken together, these lines of evidence indicate that the degree of off-target reduction by the variants is dependent on the position and nucleotide context of the mismatches.

### GuideVar facilitates optimal sgRNA selection for Cas9 variants

We next sought to predict the sequence-specific on-target efficiency and off-target effects for HiFi and LZ3 using the datasets generated in this study. Over the past years, machinelearning models have been developed and trained to predict on-target efficiency and off-target effects of WT SpCas9 and a few Cas9 variants based on a large volume of datasets (17,22,26,35). To take advantage of these pre-trained models, we developed a transferlearning framework, named GuideVar, by which previous models on WT SpCas9 and HF1 are “transferred” to new models to enhance the predictive power on HiFi and LZ3.

GuideVar consists of two modules: GuideVar-on for the prediction of on-target efficiency, and GuideVar-off for the prediction of off-target effect. In GuideVar-on, we transferred two long short-term memory (LSTM) models in DeepHF that was trained on 50,000+ sgRNAs (26). As shown in Figure 4A, the flattened layers of the LSTM models were concatenated, together with the mono- and di-nucleotide sequence features (see Methods), as the inputs of a multi-layer neural network that was trained using the HiFi and LZ3 data in this study. Based on 10-fold cross-validation, the transfer-learning model significantly improved the predictive power of on-target efficiency compared to the two source models in DeepHF and other machine-learning methods (Figure 4B). We further evaluated the performance of GuideVar-on using an independent TTISS dataset in which the on-target indel rates were measured for 59 sgRNAs with the expression of HiFi or LZ3 (18). As the result, GuideVar-on outperformed two recent deep-learning based models for the prediction of sgRNA efficiency (Figure 4C and Supplementary Table S5), suggesting its robustness for applications with HiFi and LZ3.

**Figure 4.**
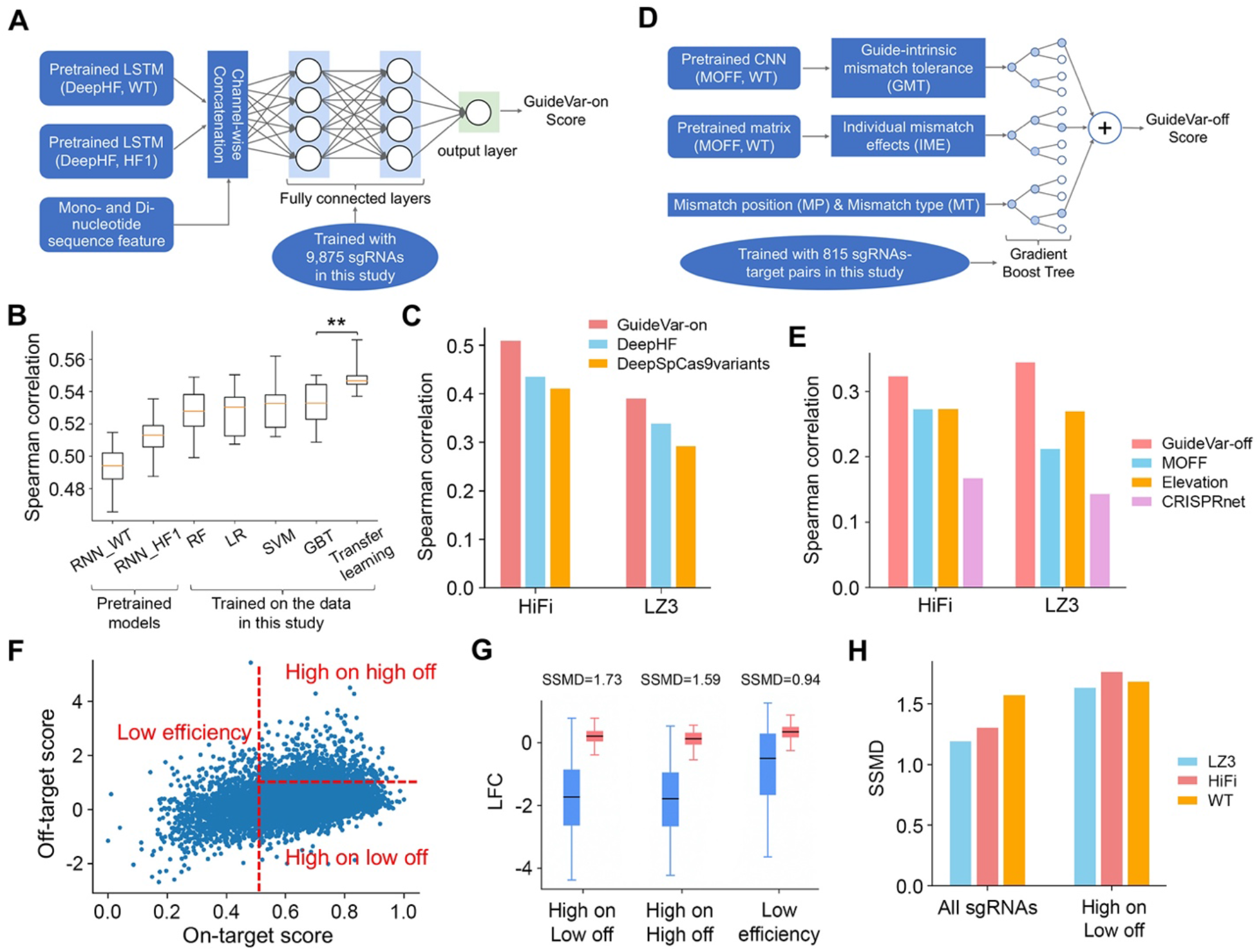
GuideVar predicts on-target efficiency and off-target effects of HiFi and LZ3 using transfer-learning. **(A)** Schematic representation of GuideVar-on for predicting on-target efficiency. **(B)** Performance comparison of different machine-learning models using 10-fold cross-validation. The performance was assessed based on the Spearman correlation between the predicted and observed efficiency scores. **, p=0.0086 (Manny-Whitney *U* test). **(C)** Performance comparison between GuideVar-on and the other two deep-learning methods for HiFi and LZ3 based on an independent TTISS dataset (18). **(D)** Schematic representation of GuideVar-off for predicting off-target effect. **(E)** Performance comparison between GuideVar-off and other off-target prediction tools for HiFi and LZ3 based on the TTISS dataset. **(F)** The distribution of all the sgRNAs in the EpiC library with on-target scores predicted by GuideVar-on and off-target scores by GuideVar-off. These sgRNAs are categorized into three groups: “Low efficiency”, “High efficiency high off-target” (high on high off) and “High efficiency low off-target” (high on low off). **(G)** Boxplot comparing the dropout effects of the sgRNAs targeting essential (blue) and nonessential (red) genes among different sgRNA categories in the screen with HiFi. Strictly standardized mean difference (SSMD) scores were computed to measure the effect size of each sgRNA category defined in (F). **(H)** Effect size of the EpiC library before and after sgRNA selection using GuideVar, in the screens with WT SpCas9, HiFi or LZ3. The effect size is measured in SSMD.

Recently, we developed MOFF, an off-target predictive model for WT SpCas9 (22). MOFF integrates multiple sequence-dependent factors, including individual mismatch effect (IME) parameterized with a matrix and a guide-intrinsic mismatch tolerance (GMT) modelled using a convolutional neural network (CNN). We note that the degree of off-target reduction by the variants is dependent on the mismatch position (MP) and mismatch type (MT) (Figure 3E and F). Therefore, GuideVar-off incorporates MP and MT, together with the quantitative measures of IME and GMT from MOFF, as the inputs for model learning (Figure 4D). Based on cross-validation, we tested a panel of linear or nonlinear machine-learning models, including linear regression (LR), random forest (RF), support vector machine (SVM), and gradient boost tree (GBT). Among them, GBT achieved the best performance (Supplementary Figure S6), thus it was implemented in GuideVar-off. We observed progressive improvement in predictive power when the four types of sequence features (GMT, IME, MT, and MP) were successively added to the model (Supplementary Figure S7), suggesting that all the features are associated with off-target effects of the variants. To predict off-target effects when multiple mismatches are present, GuideVar-off adopted the δ coefficients in MOFF to compute the combinatorial effect of multiple mismatches (22). We compared the performance of GuideVar-off to other state-of-art methods based on the TTISS dataset (18), where GuideVar-off consistently outperformed other models for both HiFi and LZ3 (Figure 4E and Supplementary Table S6).

Finally, we built a unified computational framework that integrates the GuideVar-on and GuideVar-off for the selection of optimal sgRNAs in applications with HiFi and LZ3. To test the framework, we applied GuideVar to all the sgRNAs in the EpiC library (Figure 4F), resulting in 7,553 (61.8%) sgRNAs that were predicted to be of high efficiency and low off-target effects, as well as 681 (5.6%) highly off-targeting sgRNAs and 3,980 (32.6%) inefficient sgRNAs. For each category of sgRNAs, we compared the dropout effects of the sgRNAs that target essential genes and nonessential genes and calculated strictly standardized mean difference (SSMD), a measure of the effect size of sgRNA libraries (36–38). As expected, the sgRNAs in the high-efficiency low-off-target group showed the highest effect size (SSMD=1.73 and 1.64 for HiFi and LZ3, respectively) (Figure 4G and Supplementary Figure S8). The high-fidelity Cas9 variants are rarely used in high-throughput CRISPR screens due to the compromise of efficiency. Consistently, we observed lower SSMD scores for HiFi and LZ3 when the entire EpiC library was used for screen. However, with the aid of GuideVar, the selected subset of sgRNAs showed almost equivalent effect size among WT SpCas9, HiFi and LZ3 (Figure 4H). Therefore, GuideVar will potentially facilitate the usage of high-fidelity Cas9 variants for high-throughput CRISPR screens when off-target effect is a critical concern.

## Discussion

High-fidelity Cas9 variants have been developed to reduce the off-target effect of CRISPR/Cas9 system, but the variants could also compromise the on-target efficiency which limits their applications. In this study, we found that the efficiency loss of HiFi and LZ3 is guide-specific and dependent on the sequence of the spacer. Importantly, the sequence context at the positions 15-18 relative to PAM is highly associated with the efficiency loss. Because nucleotides at positions 15-18 of the RNA-DNA heteroduplex interact with the REC3 domain of Cas9 where mutations are introduced to the variants, our results suggest a variant-specific sequence rule for on-target efficiency. Of note, most Cas9 variants developed to date harbor mutations in the REC3 domain (12,13,28). We observed similar sequence rules on HiFi and LZ3 to HF1, another REC3-mutant variant, using a dataset generated from an orthogonal study (Figure 2C). Therefore, the sequence rules derived from this study are likely to be applicable to other Cas9 variants. Despite the elaborate engineering of Cas9 variants, we also observed various degrees of off-target reduction by the variants over a large panel of sgRNAs and targets. The guide-specific off-target reduction is dependent on the position and nucleotide context of the mismatch. Particularly, a mismatch at positions 15-18 is associated with effective off-target reduction by the variants, suggesting synergistic effects between REC3 mutations and those RNA-DNA mismatches at positions 15-18. In future study, it would be interesting to explore the structural conformations with REC3 mutations and the RNA-DNA mismatches for in-depth understanding of the mechanism underlying the synergistic effects.

Given the observations of guide-specific loss of efficiency and off-target reduction by the variants, it is expected that the high-fidelity variants could effectively reduce off-target effects with little sacrifice of efficiency for a subset of sgRNAs. We developed GuideVar to facilitate guide prioritization in applications with HiFi and LZ3. In GuideVar, we introduced a transfer-learning framework by which previous machine-learning models are transferred to the new model to enhance the predictive power. We demonstrated the utilization of GuideVar via a high-throughput screen, in which a computationally selected subset of sgRNAs exhibited improved performance with the Cas9 variants, where the effect size is equivalent to or even better than WT SpCas9 (Figure 4H). Therefore, we anticipate that GuideVar holds potential to broaden the application of the high-fidelity Cas9 variants in scientific and clinical settings. Moreover, the strategy of transfer-learning seems to be a useful method for future development of prediction tools for sgRNA optimization, considering the rapidly growing body of CRISPR datasets and computational models.

## Materials and Methods

### Cell culture

DLD-1 (CCL-221), H2171 (CRL-5929) and HEK293T (CRL-3216) cell lines were purchased from ATCC. DLD-1 and HEK293T cells were maintained in RPMI1640 and DMEM medium (Gibco), respectively, supplemented with 10% FBS (Sigma), 1% penicillin-streptomycin (Gibco) at 37°C with 5% CO_2_. H2171 cells were cultured in HITES medium, supplemented with 5% FBS at 37°C with 5% CO_2_. All the cell lines were regularly tested mycoplasma free.

### Plasmid construction

WT-Cas9 expression plasmid lentiCas9-Puro was created by replacing *Blasticidin* (Blast) selection gene with *Puromycin* (Puro) gene from plasmid lentiCas9-Blast (Addgene, #52962). HiFi-Cas9 and LZ3-Cas9 constitutive expression plasmids lentiHiFi-Puro and lentiLZ3-Puro were constructed by the Gibson Assembly Site-Directed Mutagenesis approach (NEB, #E2621) using lentiCas9-Puro as the template. Inducible pCW-HiFi-Cas9 and pCW-LZ3-Cas9 plasmids were generated by the Gibson Assembly Site-Directed Mutagenesis from pCW-Cas9 (Addgene, #50661). For endogenous gene (*YAP1* and *MTAP*) knockout, the sgRNAs were constructed into lentiGuide-Blast vector according to the protocol from Feng Zhang’s lab. The lentiGuide-Blast plasmid was generated by swapping the Blast gene into lentiGuide-Puro (Addgene, #52963). The sgRNA sequences of *YAP1* and *MTAP*: 5’-TGCCCCAGACCGTGCCCATG-3’ and 5’-TCTGCCCGGGAGCTAAAACG-3’. A sgRNA (5’-GCTTACGATGGAGCCAGAG-3’) targeting the *AAVS1* gene was used as the control.

### Lentivirus production and titration

For virus packaging, HEK293T cells (4×10^6^) were seeded into a 10-cm tissue culture dish containing 10 ml fresh medium and kept at 37°C overnight. Endotoxin-free lentiviral plasmid (lentiCas9, lentiHiFi or lentiLZ3) 4 μg, psPAX2 (Addgene, #12260) 4 μg, and pMD2.G (Addgene, #12259) 2 μg were mixed in 500 μl Opti-MEM (Gibco) with 30 μl X-tremeGene HP DNA transfection reagent (Roche, #06366236001) at room temperature for 10 min, and then dropwise added to the 10-cm dish with HEK293T cells. The supernatant containing lentivirus was filtered through a 0.45-μM syringe filter 48 h after transfection. Lentivirus was aliquoted and frozen at −80°C until use. To test endogenous gene knockout, lentivirus production was performed using the same procedure using the endotoxin-free lentiviral plasmid lentiGuide-sgYAP1 or -sgMTAP.

For virus titration, cells were seeded into 24-well plates with 1×10^5^ cells/well for 6-8 h for DLD-1 or overnight for HEK293T before adding various volumes of lentivirus. H2171 cells were seeded into a 24-well plate with 5×10^5^ cells/well and incubated with various volumes of lentivirus just after seeding. To increase infection efficiency, 8 μg/ml polybrene (Millipore, #TR-1003-G) was supplemented. After 48 h infection, cells were subjected to selection with puromycin (2 μg/ml). Cell viability was tested using CellTiter-Glo^®^ reagent (Promega, #G7572) 3 d after selection and the virus titer was determined based on the cell survival rate.

### Stable cell line construction

For the construction of DLD-1 and H2171 stable cell lines with constitutive expression of WT SpCas9 or Cas9 variants, lentivirus infection was performed with the same procedure as virus titration mentioned above using similar virus volume for WT SpCas9, HiFi and LZ3. Virus volumes were adjusted accordingly to keep consistent Cas9 expression among WT SpCas9, HiFi and LZ3. After Puro (2 μg/ml) selection, the pooled cells with individual Cas9 were subcultured and expanded for determining Cas9 expression by Western blot. For the generation of HEK293T cell lines expressing doxycycline-inducible WT SpCas9 or Cas9 variants, cells were infected with pCW-Cas9, pCW-HiFi-Cas9 or pCW-LZ3-Cas9 lentivirus followed by Puro selection. Monoclonal HEK293T cells with inducible WT SpCas9 or Cas9 variants were produced by limiting dilution. To verify the knockout efficiency of the stable cell lines, the cells were infected with lentiviral lentiGuide-sgYAP1 or -sgMTAP. The knockout of YAP1 or MTAP in the resulting cells after Blast selection was determined by Western blot. A sgRNA targeting the *AAVS1* gene was used as the control.

### Western blot

Proteins were extracted with RIPA buffer (50 mM Tris-HCl, pH 8.0, 150 mM NaCl, 1% Triton X-100, 0.5% sodium deoxycholate, 0.1% sodium dodecyl sulphate) supplemented with 1x proteinase inhibitor cocktail (Roche, #11836153001). Proteins were separated on 4-15% gradient gels (Bio-Rad) and transferred onto 0.45-μM PVDF membranes, followed by blocking with 5% non-fat milk and incubation with primary antibody against Cas9 (CST, #14697, 1:2,000), YAP1 (CST, #14074, 1:2,000), MTAP (CST, #4158, 1:2,000) or β-actin (Sigma, #A5316, 1:10,000). After overnight incubation at 4°C, the membranes were rinsed with TBST and incubated with secondary anti-rabbit or anti-mouse antibodies at 1:10,000 for 1-2 h at room temperature. After rinsing, bands were visualized with ECL (Pierce, #32106). β-actin was used as the loading control.

### sgRNA library design

The EpiC library and T1 library were synthesized and constructed as described in our previous publications (22,25). For Tiling library design, we collected a set of essential transcription factor genes for DLD-1 cell line which encode multiple domain proteins, with particular attention on zinc finger proteins. To define the essentiality, we calculated the CRISPR score using the CRISPR knockout screen dataset in DepMap (39) and set the threshold <-0.5. These proteins are largely not well identified. We manually added several genes coding proteins with well-identified domains as the positive control genes. All the sgRNAs (20 nt) targeting exons (extent 20 bp up- and down-stream) were included as long as there is a PAM (NGG). The sgRNAs for the top 50 essential genes (4 sgRNAs per gene) in our previous CRISPR knockout screen data were chosen as positive sgRNA controls (25). The 500 sgRNAs targeting intergenic regions derived from the Sabatini dataset (40) were included as the negative sgRNA controls. The redundant sgRNAs and those targeting multiple loci were removed and yielded a final tiling library with 11,762 sgRNAs. Each sgRNA was flanked with 5’ sequence: TATCTTGTGGAAAGGACGAAACACCg and 3’ sequence: GTTTTAGAGCTAGAAATAGCAAGTTAAAAT. The “g” was added to the end of the 5’ flanking sequences if the first nucleotide of the guide sequence did not begin with a “G”. This oligo library was synthesized as a pool by Custom Array Inc. (Bothell, WA). The sequences of sgRNAs are provided in Supplementary Table S1-S3.

### Plasmid library construction and transformation

We amplified the synthesized oligo library using the following primer pair GGCTTTATATATCTTGTGGAAAGGACGAAACACCG (Forward) and CTAGCCTTATTTTAACTTGCTATTTCTAGCTCTAAAAC (Reverse) with NEBNext High-Fidelity Master Mix (NEB, #M0531) for 10 cycles. The pooled library cloning process was performed according to our previous publication (25). Briefly, the amplified oligo library was purified and ligated into BsmBI-digested LentiGuide-Blast using Gibson assembly. Eight transformation reactions were conducted with 2 μl of ligation product into each tube of electrocompetent cells (Lucigen, #60242) and plated onto eight 15-cm plates with Carbenicillin selection (50 μg/ml). After 14 h, all the colonies were collected as a pool for plasmid library extraction with Endotoxin-Free Plasmid Maxiprep (Qiagen, #12362).

### Pooled screen

The lentiviral libraries were prepared as described previously (25). Before determining the multiplicity of infection (MOI), DLD-1 and HEK293T cells were seeded into 10-cm dishes at the cell density of 6×10^6^ cells/dish in 10 ml fresh medium for 6-8 h and overnight, respectively. H2171 cells were seeded into 6-well plates with 5×10^6^ cells/well. For transduction, cells were incubated with different volumes of virus supplemented with 8 μg/ml polybrene for 36-48 h and then selected with Blast (20 μg/ml for DLD-1 and HEK293T, 10 μg/ml for H2171) for 5 d. Cell survival rate was measured by CellTiter-Glo assay and cell counting, and MOIs for different virus volumes were calculated compared with the non-Blast treated cells.

The pooled CRISPR knockout screens were performed using a similar procedure to the MOI testing mentioned in this section. DLD-1 with WT, HiFi or LZ3 were infected with the EpiC library or Tiling library. H2171 with WT, HiFi or LZ3 were infected with the EpiC library. Three independent replicates were set for each cell line with a minimum of 2×10^7^cells/replicate aiming for at least 500-fold coverage. Cells were infected with the lentiviral library at MOI about 0.3 for 36-48 h, followed by Blast selection for 5 d. The resulting cells were collected and passaged in fresh medium at the density of 1×10^6^ cells/ml for H2171 cells and 1×10^7^ cells/15cm-dish for DLD-1 cells. Cells were passaged every 2-3 d and cell number at a minimum of 500-fold coverage of the library was maintained during each passaging. After 20 d of maintenance (23 d for the DLD-1 cell lines with EpiC library), cells were collected for genomic DNA (gDNA) extraction using the Blood & Cell Culture Midi kit (Qiagen, #13343) according to the manufacturer’s instructions.

For off-target evaluation, we performed high-throughput screens in HEK293T cells with doxycycline-inducible WT SpCas9, HiFi or LZ3 using a synthetic dual-target system (22). The library T1 was designed with a sgRNA expression cassette paired with its relevant off-target and on-target sequences arranged in tandem. The screening procedure was performed as the screens in the DLD-1 cell line. Cell pellets were collected on day 9 after Cas9 induction for gDNA extraction.

### Library amplification and deep sequencing

The sgRNA inserts were obtained by two rounds of PCR amplification from gDNA. The first round was performed with primer pairs listed in Supplementary Table S7 with 16 cycles using NEBNext High Fidelity Master Mix (NEB, # M0541) for the screens with EpiC and Tiling libraries and NEBNext Ultra II Q5 Master Mix (NEB, #M0544) for the screens with T1 library. The 40 μg gRNA was used as the template in 8 PCR reactions for achieving about 500-fold coverage. The resulting products were purified with a PCR purification kit (Qiagen) and used 5 μl as the template for the second round of PCR with 12 cycles. The PCR products were gel purified, quantified by Qubit and qPCR, and deep sequenced using Nextseq500 at MDACC-Smithville Next Generation Sequencing Core. Single-end (75-bp) sequencing was conducted for EpiC and Tiling libraries, and 75-bp paired-end sequencing was used for the T1 library.

### CRISPR screen data processing

The pooled CRISPR knockout screen data were processed using MoPAC (https://sourceforge.net/projects/mopac/). Basically, the sequencing reads were first aligned to the sgRNA library and counted. The read counts were then processed with a quality-control module, a rank-weighted average algorithm for gene essentiality measurement, a normalization module for removing biases caused by different depths of selection, and the assessment of dropout effects of each sgRNA based on the normal distribution (Z-score).

For the high-throughput off-target screen data generated with a synthetic dual-target system, we adopted the same computational pipeline described in our previous study to call the read counts for five different types of indels and calculate the off-on ratios for each sgRNA-target pair (22).

### Sequence feature analysis for efficiency loss with Cas9 variants

To determine the sequence features associated with efficiency loss with HiFi and LZ3, we first extracted sgRNAs that target essential genes and classified them into three different groups based on their dropout effects in WT SpCas9 and the variant screens: the “Inefficient” group (Z-score >-3 in WT SpCas9 screens), the “WT efficient only” group (Z-score <-3 in WT SpCas9 screen and >2-fold in standard deviation difference between WT SpCas9 and the variant screens) and “Both efficient” (Z-score <-3 in WT SpCas9 screen and <2-fold in standard deviation difference between WT SpCas9 and the variant screens). As for HF1, the sgRNAs were also classified into three different groups based on the corresponding indel rates in WT SpCas9 and HF1 screens: the “Inefficient” group (indel rate <0.6 in WT SpCas9 screen), the “WT efficient only” group (indel rate >0.6 in WT SpCas9 and >2-fold in standard deviation difference between WT SpCas9 and HF1 screens) and “Both efficient” (indel rate >0.6 in WT SpCas9 screen and <2-fold in standard deviation difference between WT SpCas9 and HF1 screens). We then calculated the log odds ratios of different types of nucleotides at each position along the spacer sequence between the “Both efficient” and “WT efficient only” groups. The sequence logos were generated using the logomaker software (41).

We use the KL divergence to assess the importance of different nucleotide positions. Let 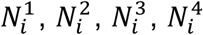 represent the frequency of A, T, C, G at the *i*th position in the “Both efficient” group and 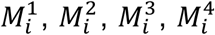 represent the frequency of A, T, C, G at the *i*th position in the “WT only” group. The KL divergence was calculated as follows:

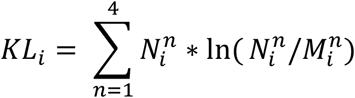

Larger KL divergence values indicate larger differences between two groups of sgRNAs at a specific nucleotide position and higher importance of the specific position for the efficiency loss with Cas9 variants.

### Efficiency prediction for HiFi and LZ3

To predict the on-target efficiency for Cas9 variants using the CRISPR screen data generated in this study, we extracted sgRNAs targeting essential genes in the variant screens with both Tiling and EpiC libraries, which resulted in 9,875 sgRNAs in the training dataset. For each sgRNA, we averaged the Z-scores in the HiFi and LZ3 screens as the final measurement for the knockout efficiency of Cas9 variants.

We first tried four conventional machine-learning methods: a linear regression model (LR), a support vector regression model with RBF kernel (SVM), a random forest regression model (RF) and a gradient boost tree regression model (GBT). All the methods were implemented through the corresponding modules from the scikit-learn package in python. Next, we implemented a two-step transfer-learning strategy to predict the on-target efficiency of sgRNAs for Cas9 variants. In the pre-training step, we took two recurrent neural network (RNN) models as our source models, which were pre-trained on the large datasets with more than 50,000 sgRNAs in WT SpCas9 and SpCas9-HF1 screens, respectively. The Bidirectional long short-term memory (BiLSTM) layers of two RNN models were then frozen to avoid destroying any of the information they contain during future training rounds. In the fine-tuning step, the outputs of BiLSTM layers of the two RNN models together with mono- and di-nucleotide sequence features were passed to downstream dense layers, of which the parameters were further fine-tuned with HiFi and LZ3 CRISPR knockout screen data described above. For the mononucleotide features, the 20-nt sgRNA sequence was binarized into a 4 × 20 two-dimension array, with 0s and 1s indicating the absence or presence of 4 different nucleotides (A, T, C, G) at every single position. For the dinucleotide features, the 20-nt gRNA sequence was binarized into a 16 × 19 two-dimension array, with 0s and 1s indicating the absence or presence of 16 different dinucleotides (AT, AC, AG, AA, TT, TA, TG, TC, CC, CA, CG, CT, GG, GA, GT, GC) at each of the constitutive positions along the sgRNA sequence.

To evaluate the performance of different methods, we adopted a cross-validation approach by randomly splitting the dataset 10 times into two groups, among which 80% were used as a training set and 20% were used for the testing. The performance was assessed by computing the Spearman correlation between the predicted and observed efficiency scores. For the independent evaluation of our transfer-learning-based model which we named GuideVar-on, we compared our model to other two deep-learning-based models DeepHF (26) and DeepSpCas9variants (17) on a dataset generated in the previous study, which includes the on-target indel rates for 59 sgRNAs targeting different genomic loci with different Cas9 variants (18). DeepHF is the combination of the RNN model and important biological features, which predicts the efficiency of sgRNAs for eSpCas9(1.1), SpCas9-HF1 and WT SpCas9. The source code of DeepHF is downloaded from https://github.com/izhangcd/DeepHF. DeepSpCas9variants is a CNN-based deep-learning model that predicts the activity of 16 different types of variants at any target sequence. The source code of the DeepSpCas9variants is downloaded from https://github.com/NahyeKim/DeepSpCas9variants. Of note, only the models predicting the efficiency for SpCas9-HF1 were used for the comparison. The performance was evaluated as the Spearman correlation between the predicted efficiency score and experimentally measured indel frequency for each sgRNA.

### Calculation of mismatch-associated energy barrier

To quantify the energy barrier associated with a specific mismatch type between sgRNA and the targeted DNA, we computed the difference of base-stacking energy between DNA-DNA and DNA-RNA hybridization over three nucleotides centered at the mismatch position based on the nucleic acid duplex energy parameters adopted from the previous study (33). To determine the influence of the energy barrier on reducing off-target effects by the variants, we compared the ratio of off-target effects (off-on ratio) between the variants and WT SpCas9 (VT/WT ratio) at each percentile of expected energy barriers for specific mismatch types. We also calculated the Pearson correlation between the expected energy barrier for a certain mismatch type and the median VT/WT ratio of all sgRNA-target pairs with that type of mismatch.

### Off-target prediction for HiFi and LZ3

To predict the sequence-specific off-target effects for HiFi and LZ3, we first collected the data for all the 1-mismatch (1-MM) sgRNA-target pairs in the high-throughput screens with HiFi and LZ3 using the dual-target system. The indel frequencies were calculated at the on-target sites and off-target sites, respectively, in the two variant screens. Considering the high correlation of both on-target rates and off-target rates between two different Cas9 variants, we averaged the on-target rates of HiFi and LZ3 and the off-target rates and then computed the averaged off-on ratios for the assessment of the off-target effects at specific target sequence for the two Cas9 variants. Finally, we got off-on ratios for 815 sgRNA-target pairs with different mismatch types and positions as our training set.

We model the sequence-specific off-target effect as the combination of four factors: (1) the individual mismatch effect (IME), which is the averaged effect for a certain type of mismatch that occurred at the specific position of the sgRNAs derived from our previous WT SpCas9 screen (22). (2) The guide-intrinsic mismatch tolerance (GMT), which measures the off-target tolerance from the intrinsic sequence of a sgRNA and is predictable with a CNN model. (3) The mismatch type (MT), which describes the type of the mismatch between a sgRNA and the target sequence. The MT is encoded as a 12-bit binary vector representing the occurrence of 12 different mismatch types rA:dA, rA:dC, rA:dG, rC:dA, rC:dC, rC:dT, rG:dA, rG:dG, rG:dT, rU:dC, rU:dG and rU:dT, where the value for the exactly occurred mismatch type is 1 and all the others are 0s. (4) The mismatch position (MP), which describes the position of the mismatch between a sgRNA and the target sequence. The MP is encoded as a 20-bit binary vector representing 20 positions along the target sequence, where the value at the mismatch-occurring position is 1 and all the other positions are 0s. We tested a simple linear regression (LR) model as well as three non-linear models including a support vector regression model with RBF kernel (SVM), a random forest regression model (RF) and a gradient boost tree regression model (GBT), to integrate four factors described above. All the methods were implemented through the corresponding modules from the scikit-learn package in python. To evaluate the performance of different methods, we adopted a cross-validation approach by randomly splitting the dataset 20 times into two groups, among which 80% were used as a training set and the remaining 20% were used for the testing. The performance was assessed by computing the Spearman correlation between the predicted off-target effects and experimentally measured off-on ratios.

Given sgRNA-target pairs with *N* (*N* > 1) mismatches, we first predict the off-target effects for each single-mismatched pair with the machine-learning models described above, denoted as *m*_1_, *m*_2_, *m*_3_, …, *m*_n_. The overall off-target effect is computed by incorporating the combinatorial effects (CE) estimated from our previous WT SpCas9 screen data. We denote *δ_ij_* as the CE between two mismatches that occur at the *i*th and the *j*th nucleotide relative to the PAM (*i*, *j* = 0, 1, 2, … , 20). The overall combinatorial effect is computed as the geometric mean of all the possible combinations among the *N* mismatches:

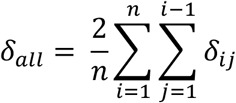

The final off-target effect (GuideVar-off) is then computed as the following formula:

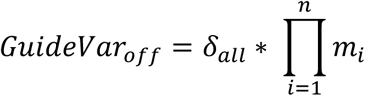

Considering that there are no off-target prediction methods specifically for Cas9 variants, we sought to compare GuideVar-off to the three state-of-art methods used to predict off-target effects for WT SpCas9: (1) Elevation is a two-layer regression model where the first layer learns to predict the effects of single-mismatch and the second layer learns how to combine single-mismatch effects into a final score (42). The source codes of Elevation are downloaded from https://github.com/Microsoft/Elevation. (2) CRISPR-Net is a more recent deep-learning method using a recurrent convolutional network, which shows superior performance compared to other machine-learning approaches (43). The source codes to implement the CRISPR-Net are downloaded from https://codeocean.com/capsule/9553651/tree/v1. (3) MOFF is a model-based off-target predictor that includes three factors corresponding to the multiplication of IME, CE and GMT (22). The source codes to implement MOFF are downloaded from https://github.com/MDhewei/MOFF. We compared the methods on the TTISS dataset, which contains 630 detected off-target sites across 59 gRNAs with WT SpCas9, HiFi, LZ3 and SpCas9-HF1 (18). For each sgRNA-target pair, we measured the off-target effect as the off-on ratio, which is calculated as the detected off-target reads divided by on-target reads. We adopted the Spearman correlations between the measured and predicted off-on ratios for quantitative evaluations.

### sgRNA design for high-fidelity Cas9 variants using GuideVar

To facilitate the rational sgRNA design for the CRISPR applications using high-fidelity Cas9 variants, we built a unified computational framework GuideVar, which consists of the following steps: First, given the sequences of a group of sgRNA candidates, GuideVar will predict the on-target score for each sgRNA using GuideVar-on. Second, GuideVar will map the sgRNAs to the genome to search for potential off-target sites harboring up to 5 mismatches using CRISPRitz, a software for rapid and high-throughput *in silico* off-target site identification (44). A final off-target score for each sgRNA is calculated as the logarithm of the sum of the predicted off-target effects for all the potential off-target sites. Third, GuideVar will classify all the sgRNAs into three categories based on the predicted on- and off-target scores: the first class is of high on-target score (>0.5) and low off-target score (<1.0), which will be selected in priority; the second class is of high on-target score but high off-target score (>1.0), which can still be used but less preferred; and the third class is of low on-target score (<0.5), which is highly recommended to discard. The source codes of GuideVar are available at https://github.com/MDhewei/GuideVar.

To test the framework, we applied GuideVar to all the sgRNAs in the EpiC library and compared the average LFCs of sgRNAs targeting essential genes and nonessential genes within each category classified by GuideVar. To quantitate the separation, we computed the strictly standardized mean difference (SSMD) between the sgRNAs targeting essential and nonessential genes. A higher SSMD indicates greater separation between the essential and nonessential genes (36).

## Data availability

The data that support the findings of this study are available from the corresponding author upon reasonable request.

## Code availability

The source codes of GuideVar are available at https://github.com/MDhewei/GuideVar.

## Acknowledgments

This work was supported by the CPRIT grant RR160097 (to H.X.) and the NIH grant R35GM137927 (to H.X.). H.X. is a CPRIT Scholar in Cancer Research.

## Author contributions

H.X., L.Z. and W.H. conceptualized the study. L.Z. and R.F. performed the experiments. W.H. performed computational analysis. H.X., W.H. and L.Z. provided data interpretation. H.X. supervised the project. All authors participated in writing the manuscript.

