## Supplementary Figures for "Guide-specific loss of efficiency and off-target reduction with Cas9 variants"

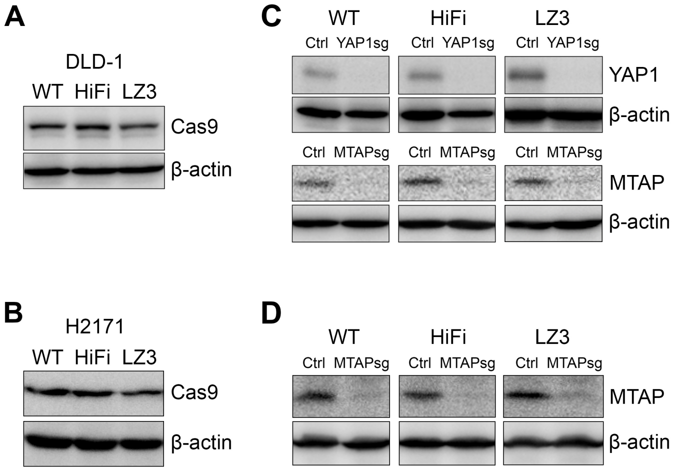


**Supplementary Figure S1. Verification of Cas9 expression and gene knockout in the engineered cell lines. (A,B)** Cas9 protein expression in **(A)** DLD-1 and **(B)** H2171 cell lines with WT SpCas9, HiFi and LZ3 determined by Western blot. (**C,D**) The knockout effects of **(C)** the DLD-1 cell lines and **(D)** H2171 cell lines with WT SpCas9, HiFi and LZ3 demonstrated by the knockout of YAP1 and/or MTAP. A sgRNA targeting *AAVS1* is used as the control.

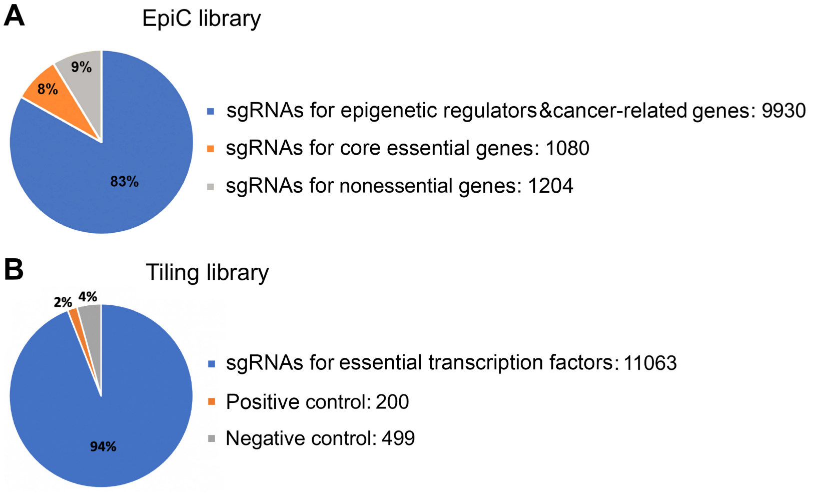


**Supplementary Figure S2. The sgRNA libraries used for CRISPR screens.** The composition of **(A)** EpiC library and **(B)** Tiling library used for the CRISPR screens with WT SpCas9, HiFi and LZ3.

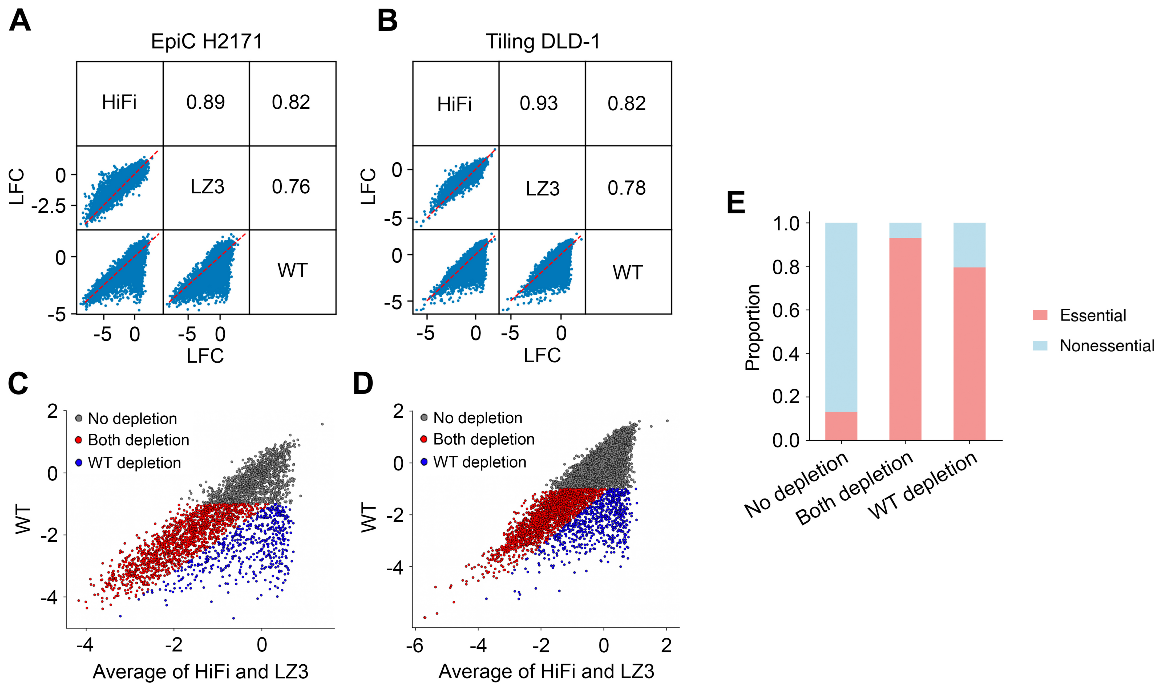


**Supplementary Figure S3. Performance of two Cas9 variants compared to WT Cas9**. **(A,B)** Pairwise comparison of the Log Fold Changes (LFCs) of sgRNAs abundance among the CRISPR knockout screens in **(A)** H2171 cell lines with EpiC library and **(B)** DLD-1 cell lines with Tiling library using WT SpCas9, HiFi and LZ3. The consistency of dropout effects was measured with the Pearson correlation. **(C,D)** Scatter plot comparing the LFC of all the sgRNAs in WT SpCas9 screens to the variant screens in **(C)** H2171 cell line with EpiC library and **(D)** DLD-1 cell line with Tiling library. Three categories were defined based on their depletion effects in WT SpCas9 and the variant screens. **(E)** The proportion of sgRNA targeting essential and nonessential genes in each category defined in **(C)**.

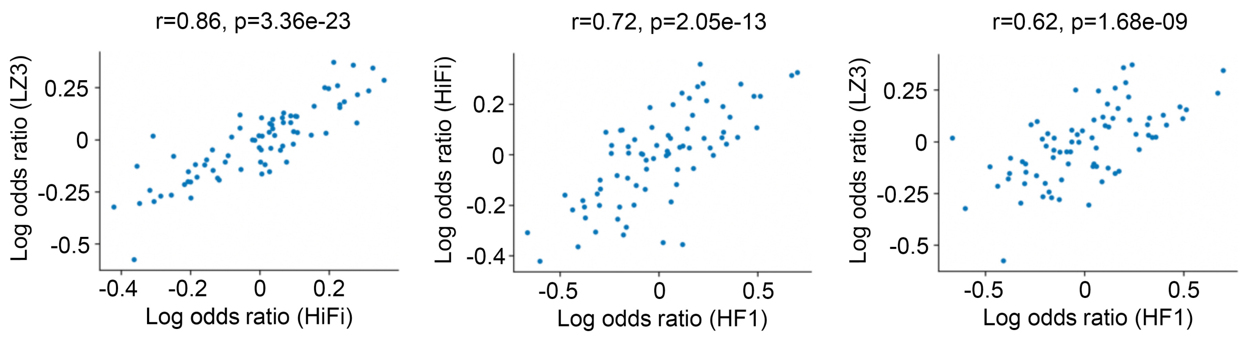


**Supplementary Figure S4. The consistency of the sgRNA sequence features identified from the screens with different Cas9 variants.** The scatter plots showing the pairwise comparison of the log odds ratios of nucleotide frequency at different nucleotide positions between “Both efficient” sgRNAs and “WT efficient only” sgRNAs among HiFi, LZ3 and HF1 variant screens.

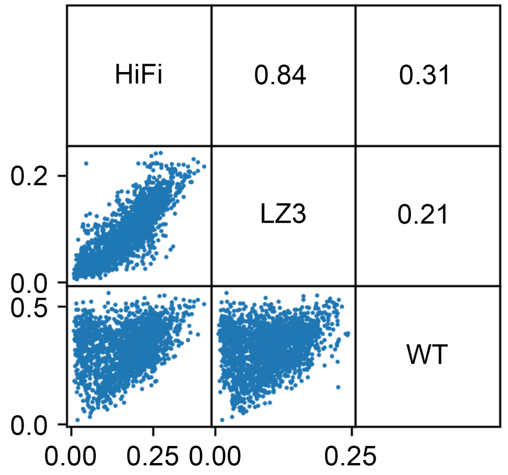


**Supplementary Figure S5. The pairwise comparison of the on-target indel rates among the CRISPR screens in HEK293T cell lines with WT SpCas9, HiFi and LZ3.** The values in the matrix indicate the Pearson correlation between different pairs.

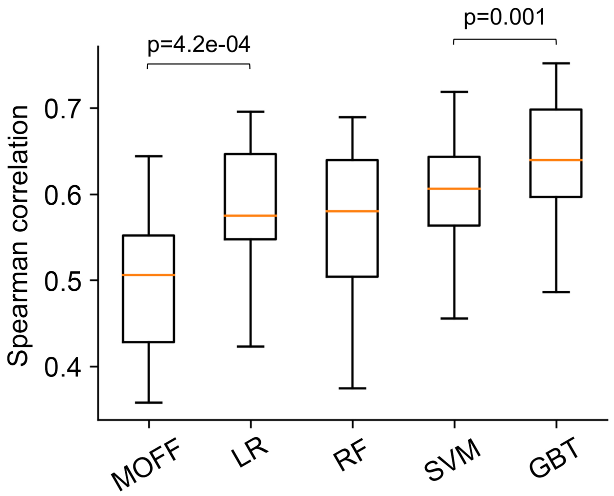


**Supplementary Figure S6. Comparison of the performance of different machine-learning models for off-target prediction.** The performance was measured by the Spearman correlation between the predicted and observed values.

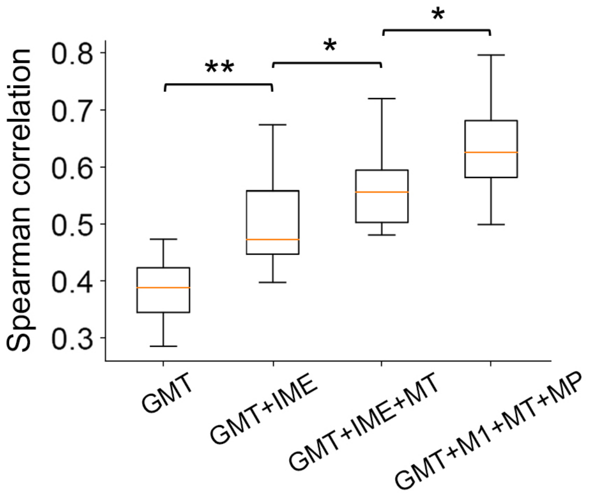


**Supplementary Figure S7. Performance comparison of GuideVar-off by successively incorporating sequence features into the model.** The performance was measured by the Spearman correlation between the predicted and observed values.

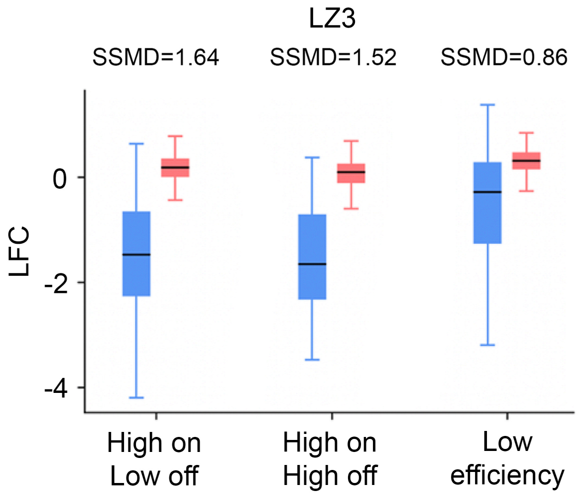


**Supplementary Figure S8. Boxplot comparing the dropout effects of the sgRNAs targeting essential (blue) and nonessential (red) genes among different sgRNA categories in LZ3 screen with EpiC library.** Strictly standardized mean difference (SSMD) scores were computed to measure the effect size of each sgRNA category.
